# Selective inhibition of the amyloid matrix of *Escherichia coli* biofilms by a bifunctional microbial metabolite

**DOI:** 10.1101/2023.07.06.547952

**Authors:** Estefanía Cordisco, María Inés Zanor, Diego M. Moreno, Diego Omar Serra

**Affiliations:** Laboratorio de Estructura y Fisiología de Biofilms Microbianos, Instituto de Biología Molecular y Celular de Rosario (IBR, CONICET-UNR), Predio CONICET Rosario, Ocampo y Esmeralda, (2000) Rosario, Argentina; Laboratorio de Metabolismo y Señalización en Plantas, Instituto de Biología Molecular y Celular de Rosario (IBR, CONICET-UNR), Predio CONICET Rosario, Ocampo y Esmeralda, (2000) Rosario, Argentina; Instituto de Química Rosario (IQUIR, CONICET-UNR), Predio CONICET Rosario, Ocampo y Esmeralda, (2000) Rosario, Argentina. Facultad de Ciencias Bioquímicas y Farmacéuticas, Suipacha 531, (2000) Rosario, Argentina

**Keywords:** Biofilms, amyloid fibers, curli, *Escherichia coli*, anti-matrix metabolites, bacillaene

## Abstract

The propensity of bacteria to grow collectively in communities known as biofilms and their ability to overcome clinical treatments in this condition has become a major medical problem, emphasizing the need for anti-biofilm strategies. Antagonistic microbial interactions have extensively served as searching platforms for antibiotics, but their potential as sources for anti-biofilm compounds has barely been exploited. By screening for microorganisms that in agar-set pairwise interactions could antagonize *Escherichia coli’s* ability to form macrocolony biofilms, we found that the soil bacterium *Bacillus subtilis* strongly inhibits the synthesis of amyloid fibers –known as curli-, which are the primary extracellular matrix (ECM) components of *E. coli* biofilms. We identified bacillaene, a *B. subtilis* hybrid non-ribosomal peptide/polyketide metabolite, previously described as a bacteriostatic antibiotic, as the effector molecule. We found that bacillaene combines both antibiotic and anti-curli functions in a concentration-dependent order that potentiates the ecological competitiveness of *B. subtilis*, highlighting bacillaene as a metabolite naturally optimized for microbial inhibition. Our studies revealed that bacillaene inhibits curli by directly impeding the assembly of the CsgB and CsgA curli subunits into amyloid fibers. Moreover, we found that curli inhibition occurs despite *E. coli* attempts to reinforce its protective ECM by inducing curli genes via a RpoS-mediated competition sensing response trigged by the threatening presence of *B. subtilis*. Overall, our findings illustrate the relevance of exploring microbial interactions not only for finding compounds with novel and unique activities, but for uncovering additional functions of compounds previously categorized as antibiotics.

**IMPORTANCE:** While traditionally serving as sources for novel antibiotics, microbial interactions have a great potential –yet to be more intensely exploited- as sources for compounds with anti-biofilm activities among other functions. Exploring such potential, we uncovered an anti-curli amyloid activity of bacillaene, a *B. subtilis* secondary metabolite, that prevents *E. coli* biofilm morphogenesis. We demonstrated that bacillaene inhibits curli by interfering with the assembly of curli subunits into amyloid fibers and that such inhibition occurs despite *E. coli* fights to reinforce its protective amyloid matrix. Moreover, we showed that bacillaene combines this anti-curli activity with a previously assigned antibiotic activity in a concentration-dependent order that potentiates the inhibitory effect against curli-based *E. coli* biofilms. The finding of additional activities of compounds previously characterized as antibiotics, as here demonstrated for bacillaene, is relevant to understand both the actual roles of secondary metabolites in modulating microbial interactions in natural niches and the potential implications of the combined activities in therapeutic applications to treat bacterial infections.

## INTRODUCTION

Within self-built communities, known as biofilms, bacteria tolerate multiple abiotic and biotic stresses. In medical contexts, this translates into biofilm-associated infections that are highly resistant to antibiotic treatments and resilient against host immune systems (1). Consequently, these infections become chronic and difficult to eradicate, posing a serious public health problem. This highlights the need for therapeutic agents that can inhibit bacterial biofilm formation. By interfering with the assembly of biofilm communities or promoting their disassembly, these agents can render bacteria more accessible and, hence, susceptible of being eliminated by host immune responses or by antibiotics in combined therapies.

The hallmark of biofilm formation is the production of an extracellular matrix (ECM) that holds the cells together (2). Due to its crucial roles in protecting the cells and shaping the overall biofilm structure, the ECM is assigned as a major target for anti-biofilm compounds (2, 3). EMCs composed of amyloid - fibers made up of protein subunits assembled in cross-β-sheet conformation-, have particularly emerged as attractive targets for anti-biofilm agents since they are abundant in biofilms of diverse bacterial species (4, 5). These are functional amyloids that unlike pathological amyloids evolved to fulfill dedicated biological roles such as adhesion, structural scaffolding/cementing and protection (6). Particularly well studied are curli amyloid fibers – simply known as “curli”- that in biofilms of most commensal and pathogenic *E. coli* strains are produced either as exclusive ECM components or combined with the exopolysaccharide phosphoethanolamine(pEtN)-cellulose (7-9). In biofilms, curli fibers are produced by starving cells in regions remote from the nutrient source (10). There, cells invest their limited resources to produce copious amount of fibers, which results in a highly dense network that literally encase the curli-producing cells conferring protection (10).

Structurally, curli are unbranched fibers that consist of two subunits: CsgB, which constitutes the initial short section, and CsgA, which forms the longest part of the fiber (9). Following translation in the cell cytosol, CsgB and CsgA are translocated via the SecYEG complex to the periplasm, where premature polymerization is prevented by the chaperone-like protein CsgC (11). Assisted by the periplasmic protein CsgE, CsgB and CsgA are then secreted through an outer membrane channel formed by the lipoprotein CsgG (12). Once outside, CsgB associates with the cell surface via interaction with CsgF, an accessory protein connected to CsgG (13). As CsgA is cosecreted, it interacts with CsgB, which induces nucleation and polymerization of CsgA into an amyloid fiber (14). While CsgB is essentially required for these events *in vivo*, CsgA alone is capable of self-polymerizing under different *in vitro* conditions (15). Since CsgB/CsgA polymerization occurs at the cell surface, it appears as an accessible target for inhibition.

Likewise, as a functional amyloid, the synthesis of the curli subunits is a highly regulated process that provides for additional potential targets for anti-curli strategies. Transcription of the *csgBAC* operon, which encodes the CsgB/CsgA subunits along with the chaperon-like protein CsgC, is activated by the biofilm regulator CsgD (16, 17). In turn, CsgD is encoded -together with curli accessory and secretion proteins- by the *csgDEFG* operon, whose expression depends on RpoS (σ^S^), the master regulator of stationary phase and General Stress Response (GSR) (10, 17). In biofilms, *E. coli* expresses RpoS mainly in response to nutrient limitation, but several other internal and external stresses can also trigger its expression (17).

Since the discovery of penicillin, antagonistic interactions among soil-dwelling microorganisms (especially *Actinobacteria*) and clinically relevant bacterial pathogens have served as a major platform for the discovery of antibiotics (18, 19). With most research attention being focused on the search for novel antibiotics (18), microbial interactions have been mostly overlooked regarding their potential as source for compounds that, rather than killing the bacteria, modulate or interfere with other bacterial behaviors such as biofilm formation. Thus, microbial interactions with focus on biofilms can allow not only to identify new compounds that specifically act on biofilm formation, but also re-evaluate metabolites previously identified as antibiotics, which, however, could also act as biofilm modulators (20).

Aiming at searching for anti-curli agents, we explored pairwise interactions between macrocolony biofilms of *E. coli* W3110, a strain that produces curli fibers as exclusive biofilm ECM components, and of soil-dwelling microorganisms, producers of secondary metabolites. We found that *B. subtilis* NCIB 3610 (abbreviated as 3610) strongly inhibits the production of curli amyloid fibers and hence impairs morphogenesis of *E. coli* W3110 biofilms. This inhibitory action on curli was found to be mediated by bacillaene, a hybrid non-ribosomal peptide (NRP)/polyketide (PK) metabolite first reported as a bacteriostatic antibiotic (21) and more recently shown to also affect biofilm formation by certain bacterial species (22, 23). We found that bacillaene combines both antibiotic and the novel anti-curli functions in a concentration-dependent manner, highlighting bacillaene as a metabolite naturally optimized for microbial inhibition. Our results reveal that bacillaene inhibits curli by directly impeding the assembly of the CsgB/CsgA subunits into amyloid fibers. Interestingly, such inhibition occurs despite *E. coli* attempts to counterbalance the inhibitory effect and to protect the cells by inducing curli genes via a RpoS-driven competition sensing response.

## RESULTS

### B. subtilis impairs morphogenesis of E. coli macrocolony biofilms by inhibiting curli amyloid fiber production

In agar-grown macrocolony biofilms of *E. coli* K-12 strain W3110, curli fibers are produced as the exclusive ECM component and are responsible for the emergence of a distinctive morphological pattern of concentric rings that is accompanied by intense red staining in the presence of the amyloid dye Congo Red (CR) (Fig. 1A). Thus, these morphological and staining patterns serve as direct readout for curli production and thereby for biofilm formation. To search for microbial compounds able to inhibit curli fibers, we screened pairwise interactions between *E. coli* W3110 and microorganisms potentially capable of producing extracellular metabolites by growing them as neighboring macrocolonies on salt-free LB agar supplemented with CR. Using this approach, we found that the soil bacterium *B. subtilis* strain 3610 exerts a strong inhibitory effect on curli production in *E. coli* W3110 macrocolonies (Fig. 1B and movie Mov. S1). Curli inhibition was readily evidenced by the absence of morphological development and CR staining in the *E. coli* macrocolony at the zone of interaction with *B. subtilis* (Fig. 1B and Mov. S1). This inhibitory effect was confirmed by microscopic analysis of cross-sections of *E. coli* W3110 macrocolonies grown close to *B. subtilis* in the presence of the fluorescent curli dye Thioflavine S (TS), which showed a large region of the upper macrocolony layer at the interaction zone being completely devoid of TS-stained curli fibers (Fig. 1D and E). Examinations at higher resolution revealed that even in areas where TS-stained curli began to be detected (*i.e*., following the regions of complete curli inhibition), the fibers appeared dispersed and unstructured, which contrasts with the highly dense network of curli fibers that form “basket”-like structures around cells in regions remote from *B. subtilis* (Fig. 1E vs 1F). The presence of bacteria in the zone of curli inhibition indicated that those cells were able to growth despite their inability to produce curli. Indeed, the number of non-curli-producing bacteria recovered in viable conditions was similar to the number of curli-producing bacteria collected viable from regions remote from *B. subtilis* (Fig. S1A).

**Fig. 1.**
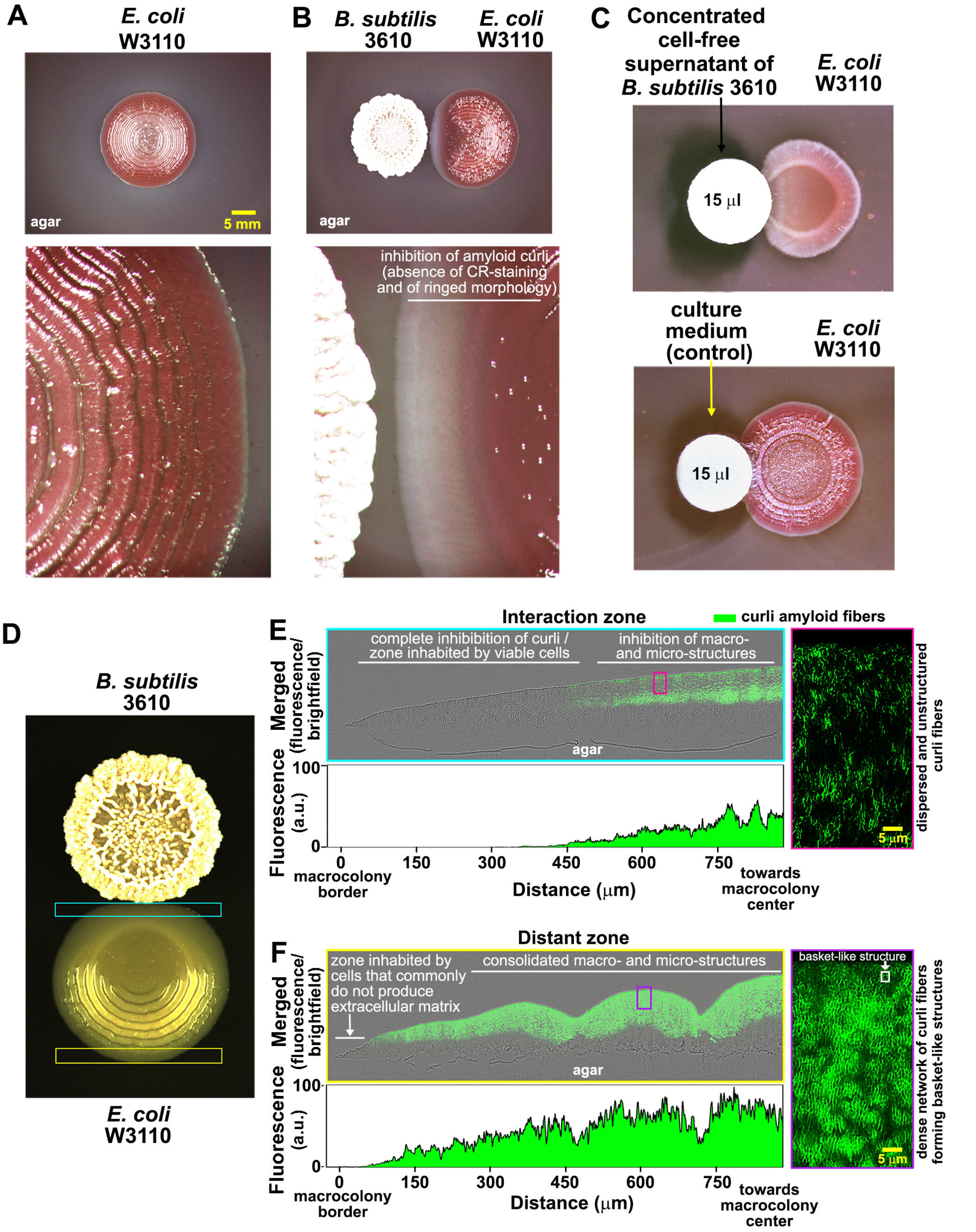
*B. subtilis* strongly inhibits curli-mediated morphogenesis of *E. coli* biofilms. Macrocolonies of *E. coli* W3110 grown on CR-containing salt-free LB agar **(A)** alone or **(B)** in close interaction with a macrocolony of *B. subtilis* 3610. The lower image in A shows at higher magnification the ringed and CR-stained patterns of a W3110 macrocolony that are dependent on curli. The lower image in B shows the absence of morphological development and CR staining in the *E. coli* macrocolony at the zone of interaction with *B. subtilis*. **(C)** W3110 macrocolonies grown in close proximity to paper discs containing concentrated cell-free supernatant (CCFS) of *B. subtilis* 3610 or culture medium (control). CCFS inhibits curli-dependent morphology and CR-staining patterns. **(D)** W3110 macrocolony grown on TS-containing salt-free LB agar in interaction with *B. subtilis* 3610. Merged fluorescence and phase contrast images of representative cross-sections through the W3110 macrocolony showing **(E)** the almost complete absence of curli at the interaction zone and **(F)** the abundance of TS-stained curli at the distant zone. Images reveal how the presence or absence of curli influence the micro and macrostructures of the colonies. Images at the right side are enlarged views of color-coded areas boxed in the cross-sections that show the amount and spatial arrangement of TS-stained curli fibers in each area. The spectral plots depict quantified patterns of TS fluorescence in respective cross-sections.

Since various *E. coli* strains can produce pEtN-cellulose as additional biofilm ECM element, we asked whether *B. subtilis* also inhibits the production of this exopolysaccharide. To test this distinguishing a potential inhibition of pEtN-cellulose from that of curli, a *csgB*-deleted *E. coli* AR3110 strain (derivative of W3110) that produces pEtN-cellulose only was set in macrocolony interaction with *B. subtilis* 3610. As shown in Fig. S1B, on CR-containing salt-free LB agar the *B. subtilis* macrocolony did not noticeably alter the morphological and CR-staining patterns dependent on pEtN-cellulose on the neighboring Δ*csgB* AR3110 macrocolony, indicating that in these conditions the inhibitory action of *B. subtilis* is not extensive to pEtN-cellulose but restricted to curli. The inhibitory action of *B. subtilis* 3610 on curli was, on the other hand, found to occur in macrocolonies of other *E. coli* strains that also produce curli as the primary ECM component such as *E. coli* MC4100 and Diffusely Adhering *E. coli* (DAEC) strain 2787 (Fig. S1C).

### The Sfp-dependent secondary metabolite bacillaene is responsible for the inhibitory effect of B. subtilis on curli amyloid fiber production

The observation that curli inhibition occurs without direct contact between *B. subtilis* and *E. coli* biofilms indicated that the effector should be a diffusible agent. To confirm this, concentrated cell-free supernatant (CCFS) prepared from 20-hour-old planktonic cultures of *B. subtilis* 3610 was tested against *E. coli* W3110 macrocolonies. As shown in Fig. 1C, CCFS adsorbed in a paper disc placed right next to the W3110 macrocolony reproduced the strong inhibitory effect on curli, confirming that the effector localized extracellularly.

*B. subtilis spp.* produces various extracellular secondary metabolites, in particular non-ribosomally synthesized peptides (NRPs) and polyketides (PKs) and hybrid PK/NRP compounds whose synthesis requires the phosphopantetheinyl transferase (Sfp, PPTase) at early steps (24). Thus, we thought that testing a *B. subtilis* 3610 mutant deficient in this enzyme (*sfp*) in biofilm interactions with *E. coli* W3110 could give us a first hint on whether a Sfp-dependent metabolite is involved in curli inhibition. Supporting our reasoning, the *sfp B. subtilis* 3610 mutant completely lost its ability to inhibit curli production and its derived macrocolony morphogenesis by *E. coli* W3110 (Fig. 2A and Fig. S2B).

**Fig. 2.**
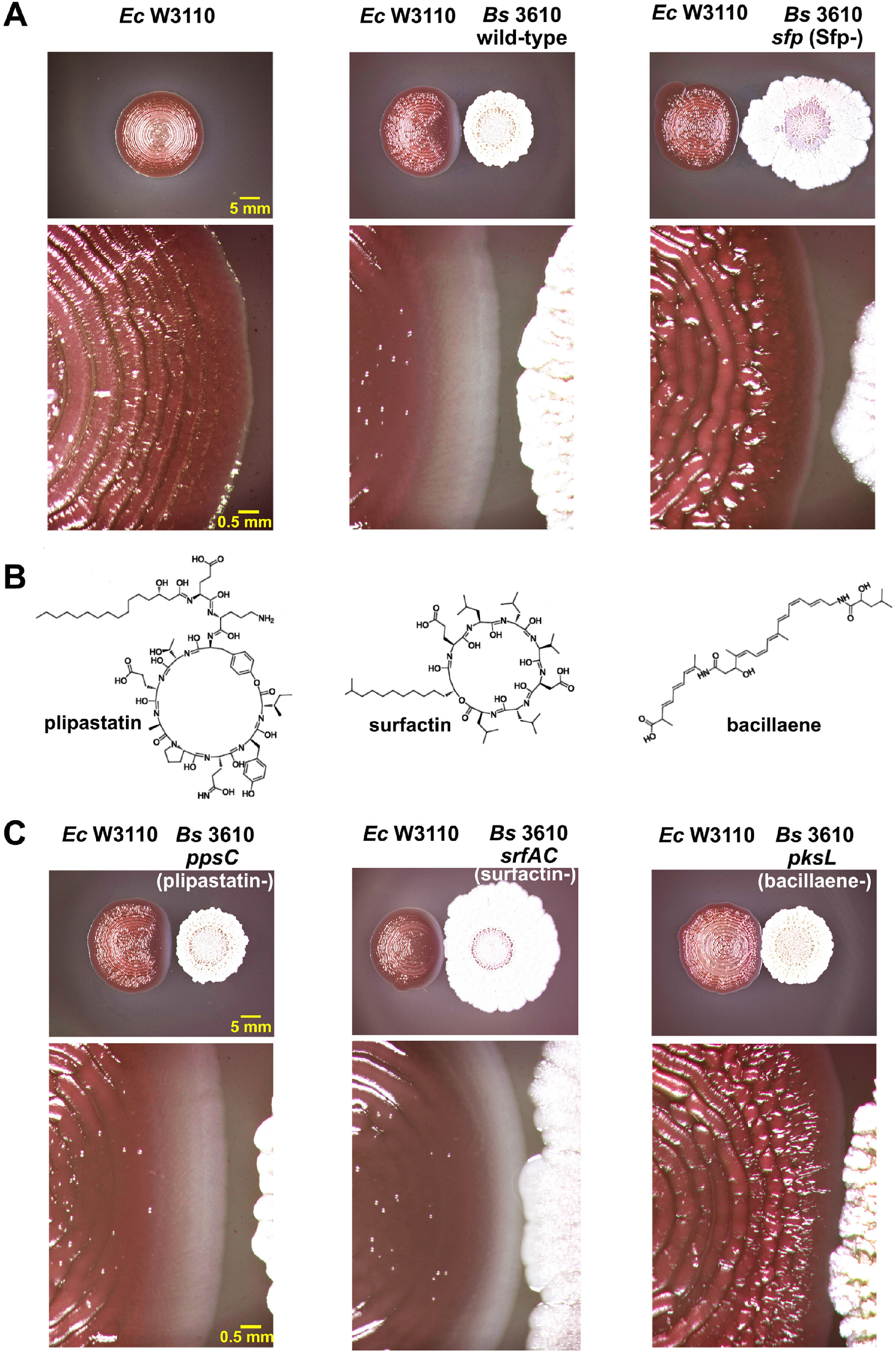
Involvement of Sfp-dependent secondary metabolites on the inhibitory effect of *B. subtilis* 3610 on curli production. **(A)** Macrocolony biofilms of *E. coli* W3110 grown on CR-containing salt-free LB agar alone or in close interaction with macrocolonies of *B. subtilis* 3610 wt or a derivative strain deficient in the phosphopantetheinyl transferase (*sfp*). The lower images are enlarged view of the *E. coli* macrocolonies shown in the upper images that magnified the zone of interaction with *B. subtilis* 3610. The images show that the lack of Sfp renders *B. subtilis* unable to inhibit curli. **(B)** Chemical structures of three Sfp-dependent secondary metabolites produced by *B. subtilis* 3610: surfactin and plipastatin, both NRPs, and bacillaene, a hybrid PK-NRP compound. **(C)** Macrocolonies of *E. coli* W3110 grown on CR-containing salt-free LB agar in close interaction with macrocolonies of *B. subtilis* 3610 strains deficient in surfactin (*srfAC*), plipastatin (*ppsC*) or bacillaene (*pksL*). The images show that in the absence of bacillaene only, curli production and hence the curli-dependent biofilm morphology occur normally.

Among the secondary metabolites whose production in *B. subtilis* 3610 requires Sfp are known surfactin and plipastatin, two cyclic lipopeptides, and bacillaene, a hybrid PK/NRP compound (24, 25) (Fig. 2B). Having these metabolites as potential effector candidates, we then tested *B. subtilis* 3610 mutants deficient in each of these compounds for their effect on *E. coli* W3110 macrocolonies. As shown in Fig. 2C, while macrocolonies of *B. subtilis* strains deficient in plipastatin (*srfAC*) or surfactin (*ppsC*) reproduced the inhibitory effect on curli, the biofilm of the *B. subtilis* strain deficient in bacillaene (*pksL*) did not affect the production of curli in the *E. coli* W3110 macrocolony, which grew developing its characteristic curli-dependent morphological and CR-staining patterns. The absence of inhibitory effect on curli by the *pksL B. subtilis* mutant was also corroborated microscopically in macrocolony cross-sections (Fig. S2E). This demonstrates that bacillaene is the Sfp-dependent metabolite responsible for the inhibition of curli.

### Bacillaene acts a bifunctional metabolite that inhibits curli-dependent formation of submerged E. coli biofilms at sub-bacteriostatic concentrations

Bacillaene was previously shown to be a bacteriostatic antibiotic that acts against various bacterial species, including *E. coli* (21, 26). This leads us to hypothesize that bacillaene could exerts bacteriostatic and/or anti-biofilm ECM effects on *E. coli* on a concentration-dependent manner. To examine this hypothesis, we first purified bacillaene produced by *B. subtilis* 3610 wild-type (wt) (Fig. S3) and tested for its minimal inhibitory concentration (MIC) against *E. coli* W3110 and its anti-curli effect on macrocolonies. Bacillaene showed a MIC value of 6 μg/ml for *E. coli* W3110, which is within the range of MIC values previously reported for *E. coli* strains (21). When a paper disc containing 5 μg bacillaene was placed close to a freshly spotted *E. coli* culture, the macrocolony biofilm grew from the inoculation point with no (or minimal) curli production until a zone close to the paper disc where it did not grow (Fig. 3A). This indicates that at high concentrations bacillaene inhibits growth, whereas at lower concentrations it inhibits curli production only.

**Fig. 3.**
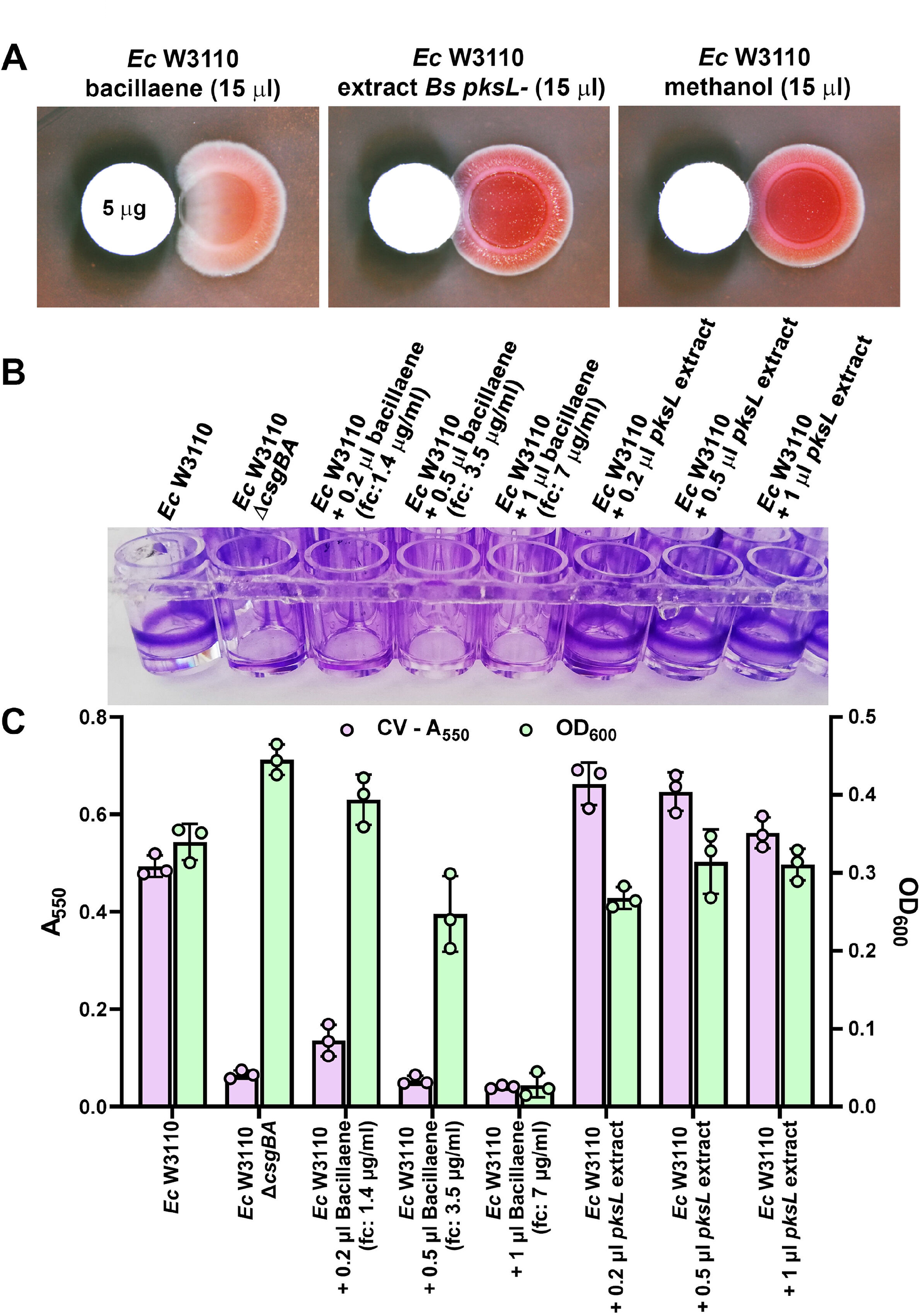
Concentration-dependent antibiotic and anti-curli effects of bacillaene. **(A)** W3110 macrocolonies grown in close proximity to paper discs containing purified bacillaene, extract from the *pksL* mutant (control) or methanol (solvent control). Bacillaene shows inhibitory effects on growth or curli production on a distance scale that reflects a differential action of the metabolite depending on its concentration. **(B)** Effects of bacillaene or *pksL* extract on the formation of curli-dependent biofilms of *E. coli* W3110 on microtiter wells. Volumes indicated correspond to a 700 μg/ml solution of purified bacillaene or to *pksL* extract. Bacillaene concentrations indicated in parentheses are final concentrations (fc) in wells. The image, which shows the presence or absence of biofilms on the well walls after CV staining, reveals that bacillaene also inhibits -at different concentration- the formation of submerged biofilms. **(C)** Quantification of *E. coli* biofilm formation and associated planktonic growth in each well determined by measuring the absorbance (A_550_) of solubilized CV and optical density (OD_578_), respectively. Data shows that at high concentrations bacillaene inhibits growth, whereas at lower concentrations it inhibits curli synthesis.

To further examine the bacteriostatic and the anti-curli effects of bacillaene, we used a multiwell-plate-based model of submerged biofilm where the sessile growth is quantified by CV staining and the planktonic growth is determined by measuring the OD_578_. In multiwell plates, *E. coli* W3110 typically forms robust biofilms on the well walls at the air-liquid interface, which is strictly dependent on curli (Fig. 3B and C). In the presence of sub-MIC concentrations of bacillaene (1.4 and 3.5 μg/ml) *E. coli* W3110 was unable to form biofilms, but able to grow planktonically (Fig. 3B and C). However, at a concentration of 7 μg/ml, *i.e.,* above the MIC, bacillaene directly inhibited both the planktonic and the biofilm growth of *E. coli.* In the presence of extract from the *pksL B. subtilis* mutant -processed in the same way as the wt extract to purify bacillaene-, *E. coli* W3110 formed biofilms and also grew in suspension (Fig. 3B and C). Altogether, these results show that bacillaene serves as a bifunctional metabolite against *E. coli*, which at high concentrations acts as an antibiotic, whereas at sub-MIC concentrations acts as an anti-curli agent preventing biofilm formation.

### Despite bacillaene inhibition of amyloid fiber production, curli genes are upregulated due to RpoS-mediated response to competition

Two mutually non-exclusive mechanisms could explain the strong anti-curli effect of bacillaene: (i) the metabolite affects expression of the curli subunits, and/or (ii) the metabolite interferes with the extracellular assembly of the subunits into amyloid fibers. To test the first scenario, we analyzed the expression of the *csgBAC* curli structural operon by setting up macrocolony interactions of *E. coli* W3110 harboring a *csgB*::*lacZ* reporter fusion and *B. subtilis* 3610 wt or its derivative *pksL* mutant, deficient in bacillaene synthesis. Using both ONPG- and X-gal-based β-galactosidase assays we did not observe downregulation of expression of the curli structural operon in W3110 macrocolonies at the zone of interaction with *B. subtilis* 3610 wt, *i.e.*, the zone where bacillaene impairs curli production, relative to a distant zone in the *E. coli* biofilm where cells exhibit normal curli production (Fig. 4A-B). On the contrary, strikingly, an increase in the expression of the curli operon at the interaction zone, independent of bacillaene, was observed (Fig. 4A-B).

**Fig. 4.**
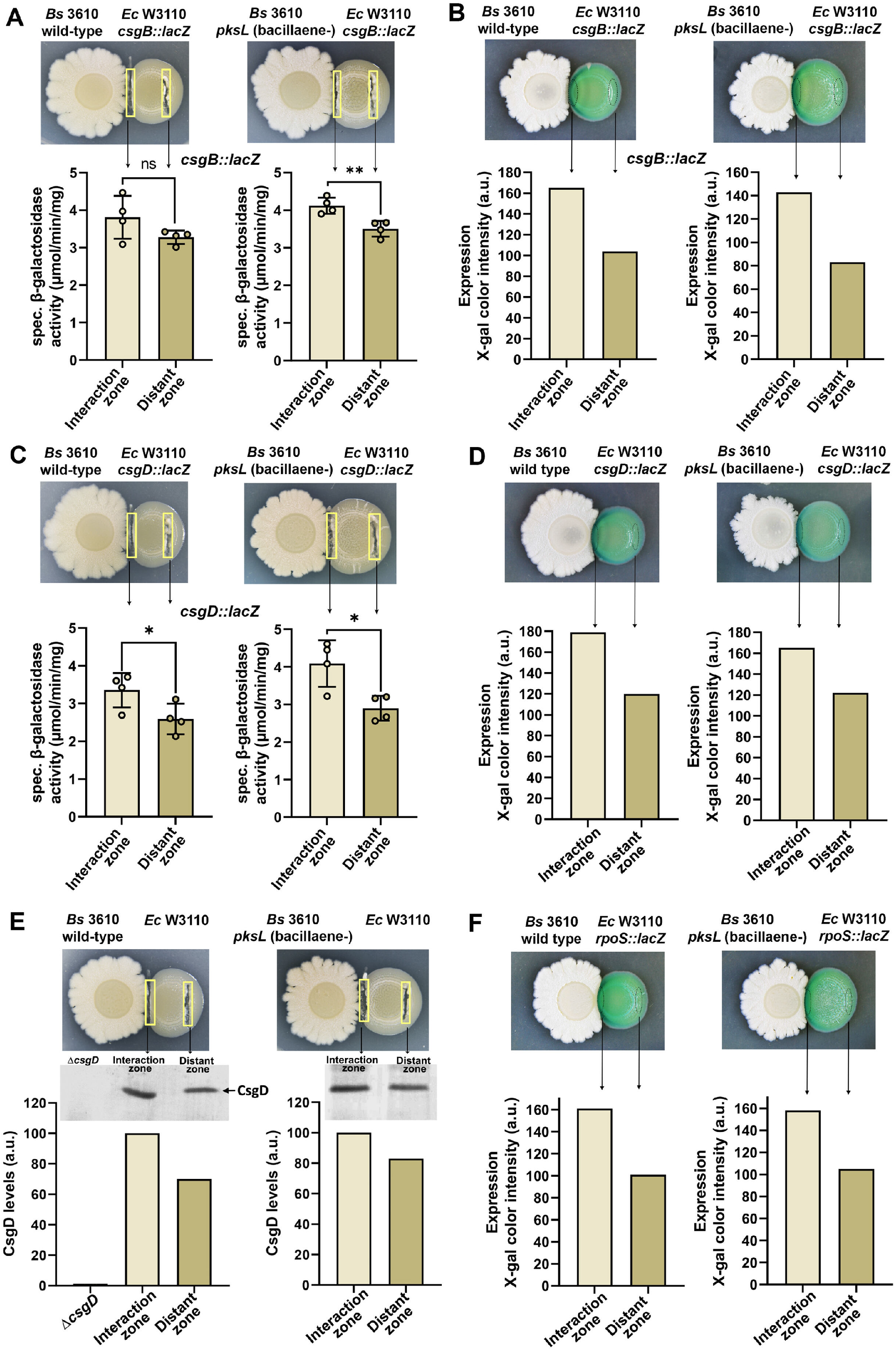
Effect of bacillaene on the expression of curli structural and regulatory genes. **(A** and **C)** Macrocolonies of *E. coli* W3110 harboring a *csgB::lacZ* **(A)** or *csgD::lacZ* **(C)** reporter fusion were grown on salt-free LB plates in close interaction with *B. subtilis* 3610 wt or its derivative *pksL* mutant. *E. coli* biomass from the interaction and distant zones -as indicated in the images- was collected and subjected to β-galactosidase activity analysis. The data are means ± standard deviations of β-galactosidase activities derived from the analysis of four independent interactions. * P<0.05; ** P<0.01; ns, no significant difference (two-tailed unpaired t-tests). **(B, D** and **F)** Macrocolonies of W3110 *csgB::lacZ* **(B)**, *csgD::lacZ* **(D)** or *rpoS::lacZ* **(F)** grown on salt-free LB plates supplemented with X-gal in close interaction with *B. subtilis* 3610 wt or its derivative *pksL* mutant. The graph depicts color intensity values of X-gal at the interaction and distant zones of respective macrocolonies shown in the figure. Data are representative of at least three independent experiments with similar results. **(E)** Cellular level of CsgD in W3110 macrocolonies interacting with *B. subtilis* 3610 wt or *pksL* mutant. *E. coli* biomass from the interaction and distant zones -as shown in the images- was collected and subjected to western blot analysis using specific antibodies against CsgD. The graph depicts the pixel intensity values of respective bands.

We extended the study to CsgD, the master regulator of biofilm formation and transcriptional activator of the *csgBAC* operon, by analyzing the expression of *csgDEFG* operon and CsgD protein levels using an *E. coli* W3110 *csgD*::*lacZ* strain and western blot, respectively. Rather than a down-regulatory effect, we observed an increase of CsgD expression -both at gene and protein levels- in the zone of interaction with both *B. subtilis* 3610 wt and *pksL* (Fig. 4C-E), which parallels the *csgB::lacZ* expression (Fig. 4A-B). This increases in *csgD/csgB* expression is independent of bacillaene and is rather due to stress induced by the presence of *B. subtilis* and/or other of its compounds at the zone of interaction, as expression of RpoS (σ^s^), the master regulator of the GSR that transcriptionally activates *csgD* and hence the *csgBAC* operon, is also significantly increased in the zone of interaction with *B. subtilis* wt and *pksL* (Fig. 4F). Remarkably, the fact that the inhibitory effect on curli production occurs when the *csgD/csgB* cascade is actually induced, indicates that the inhibitory effect exerted by bacillaene is stronger that initially observed and, on the other hand, demonstrates that *E. coli* responds to *B. subtilis* competition trying to reinforce its protective ECM via a global stress response.

These results also indicate that the inhibitory effect on curli occurs downstream *csgBAC* transcription. To confirm this, we expressed the curli structural operon from a synthetic RpoS-dependent promoter (SynP(σ^s^)::*csgBAC*) in either Δ*csgD* or Δ*csgBAC* W3110 backgrounds, to achieve production of curli amyloid fibers independent from CsgD or from the entire cascade upstream the curli operon (Fig. 5A). As shown in Fig. 5B and C, despite uncoupling the expression of the *csgBAC* operon from its natural regulation, the inhibitory effect of *B. subtilis* mediated by bacillaene persisted. This confirms that the inhibitory action of bacillaene occurs down-stream *csgBAC* transcription.

**Fig. 5.**
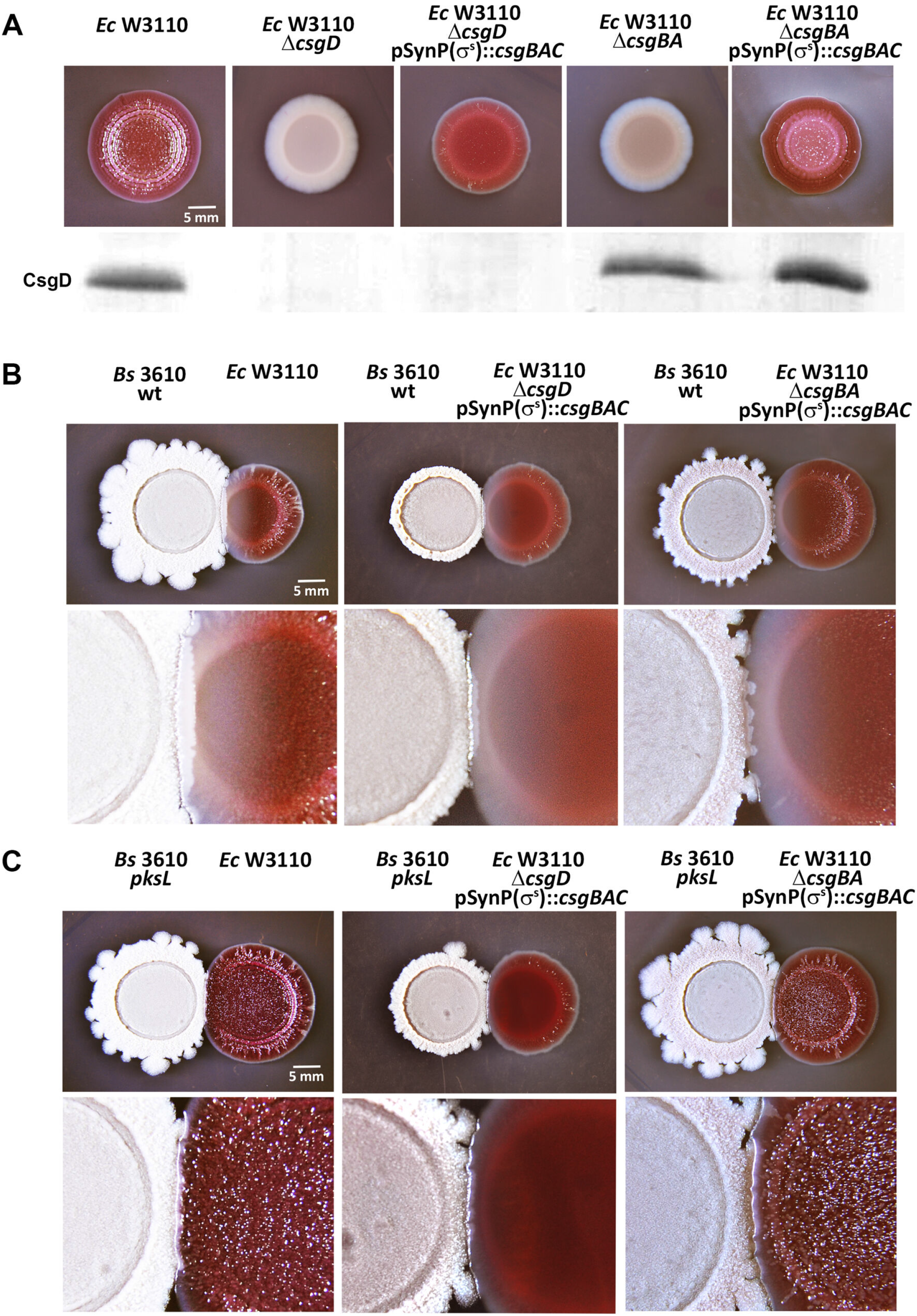
Bacillaene inhibits curli production still when expression of curli subunit genes is uncoupled from its natural regulation. **(A)** Macrocolonies of *E. coli* W3110 and derivatives Δ*csgD* and Δ*csgBA* mutants with or without expressing the *csgBAC* operon from a synthetic RpoS-dependent promoter (SynP(σ^s^)), grown on CR-containing salt-free LB agar. The western blot reveals the cellular levels of CsgD in cells from respective macrocolonies. **(B-C)** Macrocolonies of W3110 and derivatives Δ*csgD* and Δ*csgBA* mutants with or without expressing the *csgBAC* operon from the SynP(σ^s^) promoter, grown on CR-containing salt-free LB agar in close interaction with *B. subtilis* 3610 wt **(B)** or *pksL* mutant **(C)**. The lower images show at higher magnification the zone of interaction between *E. coli* and *B. subtilis* strains.

### Bacillaene impairs curli amyloid fiber assembly in vitro and in vivo

If bacillaene does not affect the expression of the curli subunits, it may alternatively affect the assembly of the subunits into amyloid fibers. In such a scenario, curli subunits would be synthesized and exported, but instead of being assembled into amyloid fibers and remain associated to the cells in the macrocolony they may disperse into the underlying agar. Thus, we first proposed to determine whether the absence of curli fibers in W3110 macrocolonies at the zone of interaction with *B. subtilis* 3610 wt correlates with the absence of CsgA, the major curli subunit. To do this, biomass of *E. coli* W3110 macrocolonies was collected at zones of interaction and distant (control) from *B. subtilis* 3610 wt or *pksL* (included as control) and analyzed for the presence of CsgA by western blot. As in control samples CsgA exists forming amyloid fibers, samples were treated with 5% SDS to efficiently depolymerizes the fibers and analyze the subunits in monomeric form. In sharp contrast to the relatively large amount of CsgA detected at a zone distant from *B. subtilis* wt, only low residual amount of CsgA was detected at the zone of interaction with *B. subtilis* (Fig. 6A). This shows a correlation between the almost complete absence of curli fibers and of curli subunits in the colony at the zone of interaction with *B. subtilis*.

**Fig. 6.**
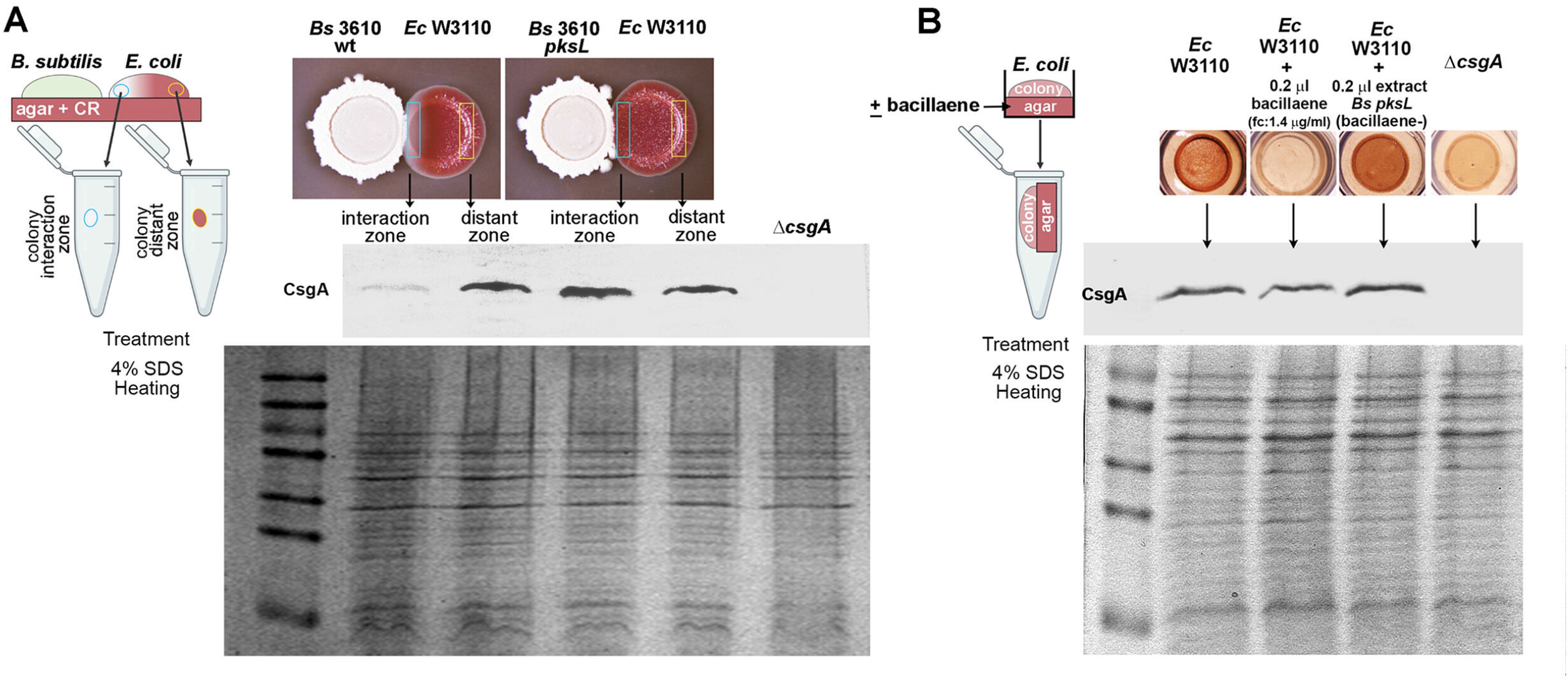
**(A)** CsgA is found in low residual amount in *E. coli* W3110 macrocolonies experiencing inhibition of curli production. Biomass at the interaction and distant zones -as indicated in the images- was collected from *E. coli* macrocolonies grown in proximity to *B. subtilis* 3610 wt or *pksL* and subjected to SDS treatment and western blot analysis to detect CsgA. The SDS-PAGE gel is shown as a sample loading control. **(B)** CsgA is detected when bacillaene-treated macrocolonies devoid of curli fibers are sampled together with the underlying agar. Colony biofilms grown in the presence/absence of bacillaene or *pksL* extract were collected with the under the underlying pieces of agar and then subjected to SDS treatment and western blot analysis for CsgA detection. The SDS-PAGE gel is shown as a sample loading control.

To investigate if, due to bacillaene effects, the curli subunits disperse into the underlying agar, *E. coli* W3110 macrocolonies were grown on agar poured in small wells with or without supplementation of bacillaene or extract from the *pksL* mutant; subsequently, each colony including the underlying piece of agar was subjected to SDS treatment and western blot analysis for CsgA detection (27, 28). Despite the bacillaene-treated macrocolony grew without evidencing CR staining, CsgA was still detected in the sample at a level comparable to that seen in samples of macrocolonies grown in the absence of bacillaene or in the presence of the *pksL* mutant extract, which exhibited CR-stained curli fibers (Fig. 6B). This shows that in the presence of bacillaene, the curli subunits accumulate in the underlying agar, pointing out that the metabolite exerts its inhibitory action at the fiber polymerization level.

To further investigate whether bacillaene affects the assembly of curli fibers, we proposed to examine the polymerization of purified CsgA *in vitro* in the presence or absence of bacillaene or extract from the *pksL* mutant. Typically, CsgA polymerization *in vitro* can be monitored using thioflavin T (ThT), a fluorophore whose fluorescence increases upon its binding to the β-sheet-rich structures of the amyloid CsgA fibers that assembles. Unfortunately, when controls for polymerization assays using ThT were set up, we found that bacillaene itself interferes with ThT fluorescence, impeding the correct monitoring of the polymerization (data not shown). Since amyloid fibers are insoluble and precipitate as they accumulate overtime and bind CR, we then took advantage of these amyloid properties as an alternative to examine the effect of bacillaene on CsgA amyloidogenesis. Thus, CsgA was purified using the expression system described by Zhou *et al* (28) and allowed to polymerize for 16 h in the presence or absence of bacillaene or extract from the *pksL* mutant. The incubation of CsgA alone in KPi buffer led to a precipitate that was confirmed by SEM microscopy to consist of a dense network of fibers (Fig. 7A and C). Such a precipitate was also observed in the presence of the *pksL* mutant extract, but it was absent in the presence of bacillaene, indicating that the metabolite prevents the accumulation of CsgA amyloid fibers (Fig. 7A). The same results were observed when the assay was performed using CR as amyloid fiber dye (Fig. 7B).

**Fig. 7.**
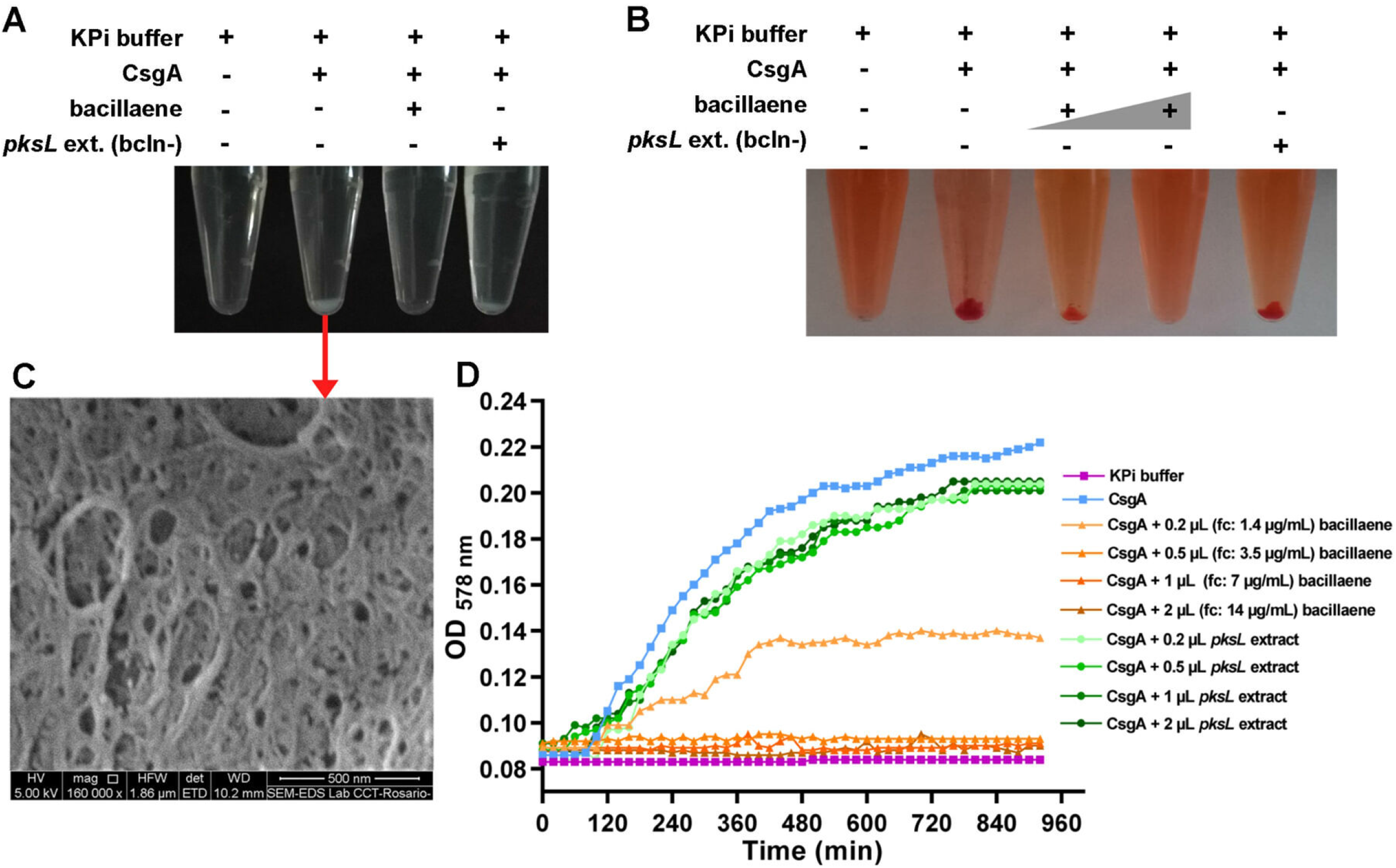
**(A-B)** Bacillaene prevents accumulation of insoluble CsgA amyloid fibers *in vitro*. Purified CsgA was allowed to polymerize for 16 h in the presence or absence of bacillaene or extract from the *pksL* mutant without **(A)** or with **(B)** supplementation of CR. CsgA polymerizes forming insoluble and CR-stainable amyloid fibers that accumulate and precipitate. Such polymerization is abolished in the presence of bacillaene. **(C)** SEM image showing the dense network of amyloid fibers that results from the polymerization of purified CsgA *in vitro* as indicated in A. **(D)** Bacillaene inhibits the polymerization of CsgA *in vitro* in a dose-dependent manner. The kinetics of accumulation of CsgA fibers in the presence/absence of bacillaene o the *pksL* extract (at different concentrations/volumes) was monitored by measuring the optical density at 578 nm. Volumes indicated correspond to a 700 μg/ml solution of purified bacillaene or to *pksL* extract. Bacillaene concentrations indicated in parentheses are final concentrations (fc) in wells. The graph shows representative data sets of at least three replicates performed for each experiment with similar results.

To gain more details, we monitored the accumulation of CsgA fibers by measuring the optical density at 578 nm. Freshly purified CsgA alone displayed a sigmoidal curve of aggregation that reflects a typical pattern of curli subunit assembly into amyloid fibers (Fig. 7D). Note that the lag phase in this *in vitro* assay is mainly due to the initial absence of an amyloid template, which *in vivo* would be provided by CsgB. While at 1.4 μg/ml bacillaene significantly diminished the accumulation of CsgA fibers, at 3.5 μg/ml it already completely abolished CsgA polymerization, showing that it is a strong inhibitor of curli fiber assembly. Consistent with this, the presence of the *pksL* extract (in different volumes) causes only minor attenuation of the CsgA polymerization pattern.

In principle, bacillaene could interact with curli subunits in their unstructured, yet unpolymerized form, or once they are folded. While predicting bacillaene binding to CsgB/A unstructured forms appears challenging, molecular docking simulations were performed to analyse the interaction between the metabolite and folded forms of the subunits. Interestingly, the lowest energy predicted position showed a putative binding site of the metabolite to CsgB in a hydrophobic groove on the upper surface of the β-helix formed by the C-terminal repeating unit R5, *i.e.*, perpendicular to the fiber axis (Fig. 8). In addition to the hydrophobic interaction between bacillaene and CsgB, the interactions of the polar groups of bacillaene with Arg147 and Gln150 are clearly noted. Such binding was also predicted for CsgA (Fig. S4), although it seems to be somewhat less strong than for CsgB due to the lack hydrophobic amino acids in that region. R5 was shown to be required for anchoring CsgB to the cell surface and to mediate -along with R1-CsgA-CsgB and CsgA-CsgA interactions (9, 29, 30). Thus, these results support a mode of action in which bacillaene impedes curli fiber assembly by preventing or conditioning such interactions.

**Fig. 8.**
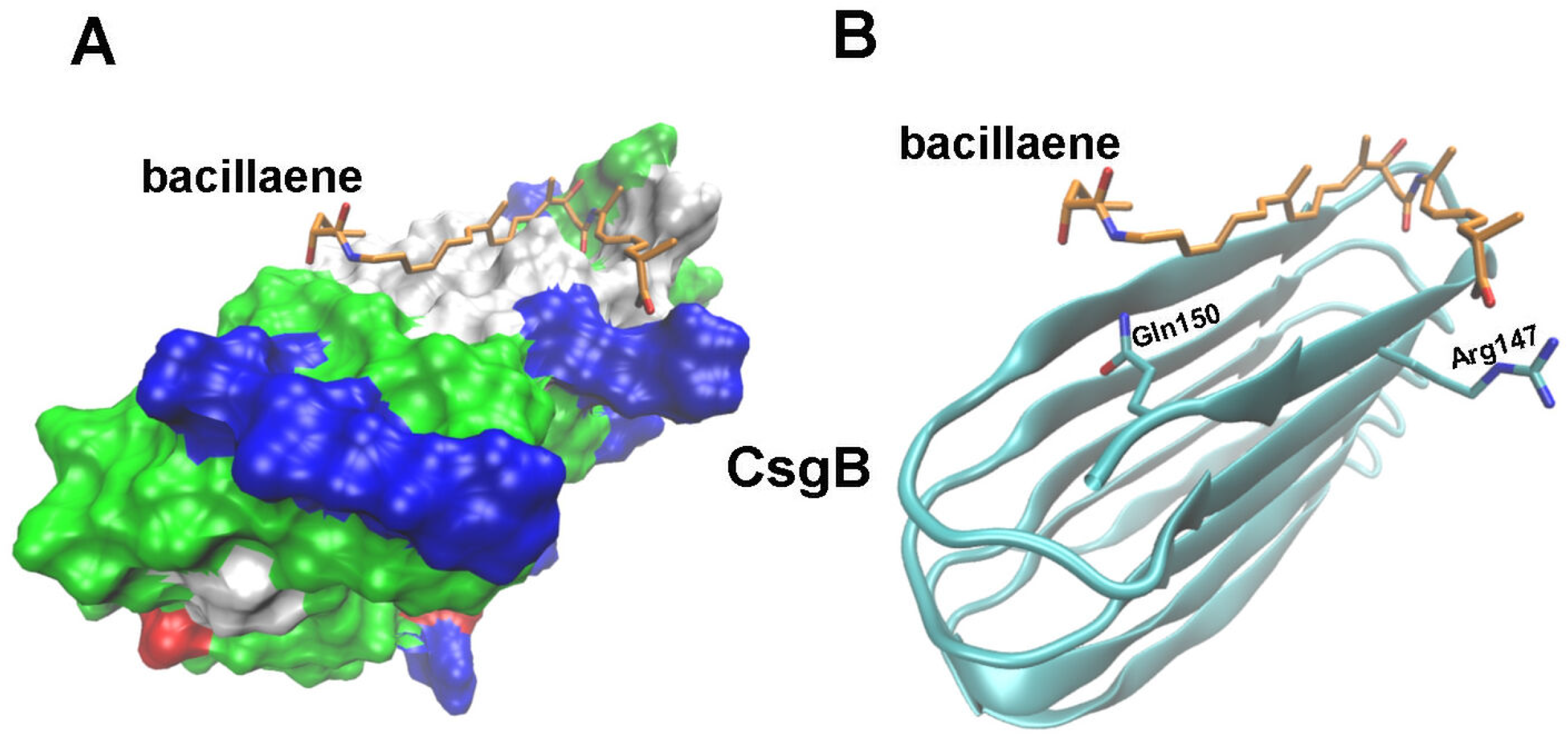
Lowest energy position of bacillaene on CsgB determined by docking simulations. **(A)** Representation of CsgB exhibiting binding of bacillaene to the surface of the β-helix formed by the C-terminal R5 repeating domain. CsgB is shown with colour coding according to the type of residues: nonpolar residues (white), basic residues (blue), acidic residues (red), and polar residues (green). **(B)** Representation of CsgB β-sheet structure highlighting the Arg147 and Gln150 residues that interact with bacillaene.

## DISCUSSION

By screening for microorganisms that upon interaction with *E. coli* in macrocolony biofilms could antagonize its ECM production, we found that *B. subtilis* strongly inhibits the synthesis of amyloid curli fibers, the primary structural elements of *E. coli* biofilms. Consequently, *E. coli* colonies lost completely their complex morphology and at the microscale cells lost their protective matrix “shield” that consists of a basket-like structure that curli fibers form around each cell. We identified bacillaene, a NRP/PK and Sfp-dependent metabolite secreted by *B. subtilis* (21, 31), as the effector molecule and could demonstrate that bacillaene exerts its inhibition by directly impeding the assembly of the CsgB/CsgA subunits into amyloid fibers. Various experimental evidences support this mode of action: (i) bacillaene does not downregulate transcription of curli structural or regulatory genes (expression of *csgBAC* and *csgDEFG* operons actually increases in cells experiencing inhibition of curli production), (ii) in the presence of bacillaene, *E. coli* cells within the macrocolony produce and secrete the curli subunits, which, however, instead of remaining associated to the biofilm cells in the form of CR-stainable amyloid fibers -as occurs in the absence of bacillaene-, diffuse away into the underlying agar, (iii) in CsgA polymerization assays *in vitro* bacillaene directly abolishes the accumulation of CsgA amyloid fibers, (iv) molecular docking simulations predict bacillaene binding to the folded form of CsgB and CsgA, particularly to a hydrophobic groove on the surface of the β-helix formed by the C-terminal R5 repeating domain, with the binding affinity for CsgB being stronger than for CsgA.

Previous studies have shown R5 to be required for anchoring CsgB to the cell surface and to mediate -along with R1-CsgA-CsgB and CsgA-CsgA interactions (9, 29, 30). Thus, by targeting such key domain is likely that bacillaene not only affects CsgB association to cells, which is necessary to efficiently template CsgA subunits into amyloid fibers *in vivo*, but also alters the interactions among subunits, directly impairing amyloidogenesis progression. Early impediment of amyloid assembly is consistent with the observation that at various bacillaene concentrations CsgA polymerization curves appear as a continuous lag phase with no further increase over time.

Predicted binding of bacillaene to the hydrophobic groove formed by repeating unit R5 suggests that the anti-amyloidogenic effect could be sequence/structure specific and hence not extensive to all bacterial amyloids. Supporting this assumption, biofilm morphogenesis by *B. subtilis* itself, which also depends on an amyloid matrix component, namely TasA (32), appeared unaffected during growth. It seems obvious that *B. subtilis* would keep its biofilm ECM untargeted from its own metabolite; unless bacillaene could serve as a self-produced biofilm disassembly factor acting in a dispersion stage. This latter, however, seems unlikely as bacillaene was shown to be produced in mature *B. subtilis* biofilms (33) and even to accelerate its structural consolidation (34).

Bacillaene was first reported as a broad-spectrum antibiotic with bacteriostatic effect (21). More recently, additional functions of bacillaene became apparent from competition studies pairing bacillaene-producing *Bacillus spp.* strains with other microorganisms, including inhibitory effects on biofilm formation by *Campylobacter jejuni* and *Salmonella Typhimurium* (22, 23). How bacillaene affects biofilm formation by these microorganisms has, however, remained unclear. Here, we show in *E. coli* that the anti-biofilm activity of bacillaene is due to its uncovered anti-curli (*i.e*., anti-ECM) effect and hence independent from its antibiotic function. While independent, both bacillaene activities operate in a concentration dependent manner, with the anti-curli/biofilm function occurring already at sub-MIC concentrations. This order of activities is likely because secreted CsgB/A subunits are readily accessible targets and thus at low concentration most bacillaene molecules would target the subunits first with a few or none molecules available to act intracellularly blocking cell growth. From an ecological perspective, this order of activities seems also advantageous for *B. subtilis* competing for space and resources against biofilm-forming *E. coli*; a “warefare” scenario where it has to fight ECM-producing cells and where distance-dependent concentration gradients of antagonistic molecules become relevant. First, bacillaene would prevent *E. coli* cells to produce their ECM protection when only low (sub-MIC) concentrations reach the community (Fig. 9A). Since ECM itself is key driver of colony biofilm expansion (35), inhibition of curli at this stage also represents a way by which *B. subtilis* can limit substrate colonization by *E. coli*. Secondly, once *E. coli* cells are unshielded and bacillaene concentration increased due to closer proximity of *B. subtilis,* it would more easily inhibit growth of proximal *E. coli* cells (Fig. 9A). The increase in bacillaene concentration at this latter stage could also result from an increase in bacillaene production induced by the proximal presence of *E. coli*, as *B. subtilis* was previously shown to upregulate bacillaene biosynthetic genes in the presence of competitors (22, 33).

**Fig. 9.**
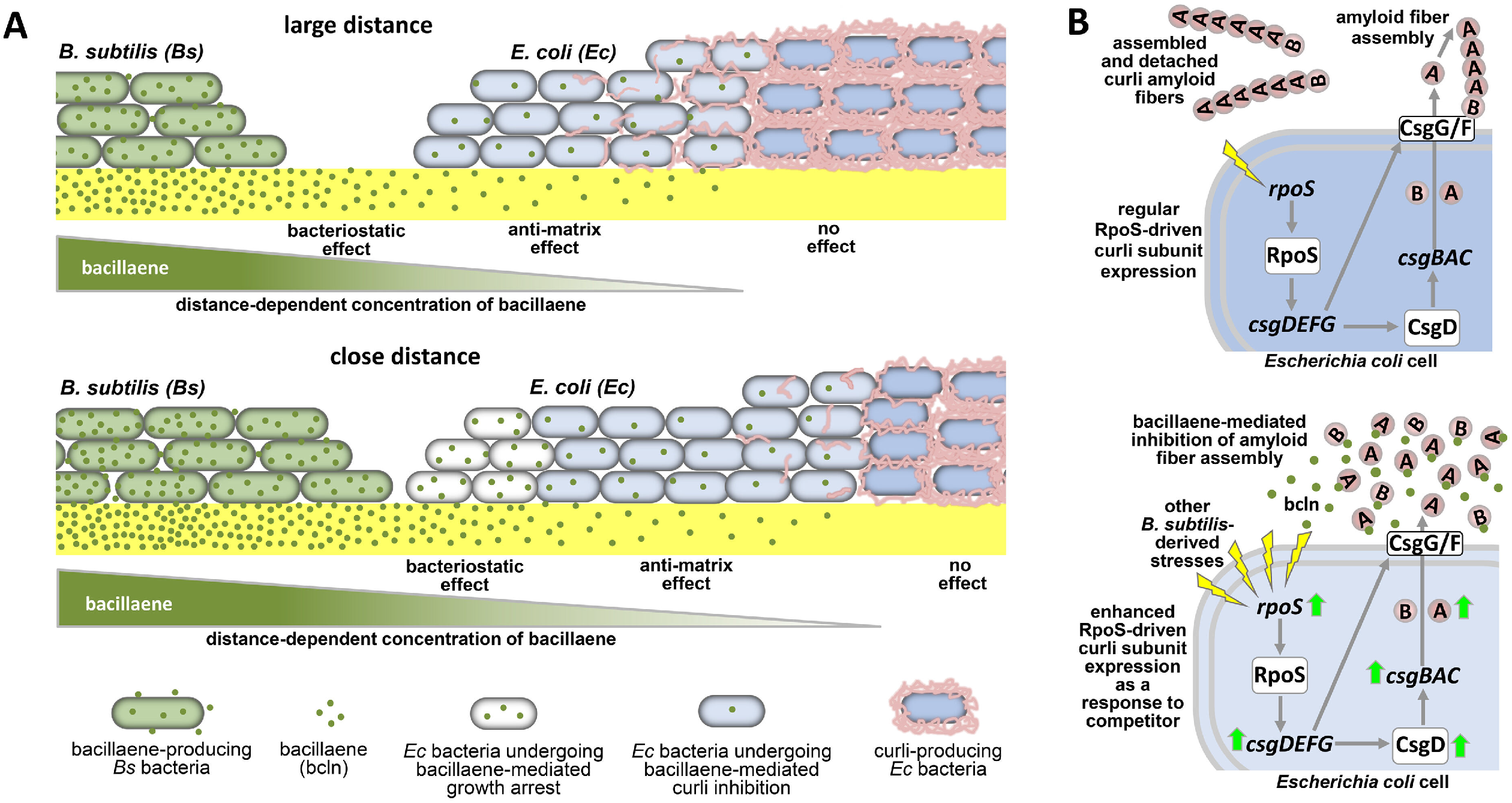
**(A)** Schematic of *B. subtilis-E. coli* colony interaction illustrating gradient-dependent inhibitory effects of bacillaene. In its competition with *E. coli* in biofilms, *B. subtilis* secretes the PK/NRP hybrid metabolite bacillaene, which diffuses away creating a distance-dependent concentration gradient. As a function of this gradient, the first bacillaene effect on *E. coli* biofilm cells is to inhibit curli assembly rendering cells “naked”, which occurs when cells are exposed to sub-bacteriostatic concentrations of bacillaene (upper panel). As *B. subtilis* and *E. coli* biofilms become closer, those *E. coli* cells that already lose their curli protection become exposed to higher bacillaene concentrations, which starts causing a second effect on *E. coli* cells, that is, to inhibit their proliferation (bacteriostatic effect) (lower panel). Thus, by combining in a single metabolite two inhibitory functions active at distinct concentrations, *B. subtilis* appears to have optimized a “weapon” to antagonize bacteria such as *E. coli*; it first renders cells devoid of their ECM protection and, secondly, once they are unshielded, it inhibits their growth. **(B)** Simplified schematic of RpoS-driven control of curli subunit expression and extracellular subunit assembly into amyloid fibers in *E. coli* cells distant from *B. subtilis*, not reached by bacillaene (upper panel). At sub-MIC concentrations, bacillaene inhibits extracellular assembly of curli subunits into amyloids fibers. *E. coli* cells react to this and other *B. subtilis* stresses trying to reinforce curli production by enhancing CsgB/A expression via a RpoS-mediated competition sensing response (lower panel).

This multifunctionality of bacillaene makes it belong to the list of antibiotics shown to alter cellular processes or bacterial behaviours –including biofilm formation- at sub-MIC concentrations (20, 36, 37). Bacillaene, however, appears to be one of the only antibiotics –together with 2,4-diacetylphloroglucinol (DAPG) produced by *Pseudomonas protegens* (38)- known to inhibit biofilm formation rather than to stimulate it, as occurs with other antibiotics (39). This highlights bacillaene as a kind of unique metabolite naturally optimized for microbial inhibition.

The finding that *E. coli* cells in close interaction with *B. subtilis* show increased expression of the *csgBAC* operon despite experiencing inhibition at the fiber assembly level, indicates that *E. coli* fights to reinforce its protective ECM (Fig. 9B). This also reflects that the anti-amyloidogenic potency of bacillaene is actually strong as it remains effective in preventing amyloid fiber formation in a context of curli subunit overproduction. Such upregulation of *csgBAC* expression is not exclusively linked to the bacillaene effect, as it still occurs in its absence. Rather, it most likely results from the contribution of other stresses associated to the threatening presence of *B. subtilis* (e.g.: nutrient depletion). This is supported by the fact that expression of RpoS is strongly induced in *E. coli* cells at the zone of interaction with *B. subtilis*. As part of its response to diverse stresses, RpoS activates transcription of the *csgDEFG* operon (10, 17). CsgD is then expressed and activates transcription of the *csgBAC* operon (10, 16). Consistent with this, CsgD was also found overexpressed in *E. coli* cells proximal to *B. subtilis*. Overall, the fact that ECM biosynthesis (and thus biofilm formation) is integrated under the control of a key global stress response and that *E. coli* cells induce such response in proximity of *B. subtilis* supports a more general hypothesis, namely competition sensing, which assumes that bacteria use stress responses to detect and respond to competitors (Fig. 9B) (40). Moreover, this *B. subtilis*-induced stress-driven increase of curli subunit expression also supports the postulate that enhancing biofilm formation is actually a common response that bacteria deploy as a way to protect themselves from competing microbes (41), even though here, in the case of *E. coli*, such enhancement was ultimately not manifested because bacillaene targeted the last step of curli production.

The encounter between *B. subtilis* and *E. coli* can naturally occur in soil, which is the habitat of *B. subtilis* and a secondary niche for many commensal and pathogenic *E. coli* strains excreted from the mammalian gut, but it can also be envisioned in the context of gastrointestinal *E. coli* infections in humans. *B. subtilis* is an admissible probiotic for humans (42), and thus its consumption could have potential implications to prevent or treat infections associated to curli-based *E. coli* biofilms. As curli amyloid fibers produced by *E. coli* and related enterics were also shown to promote human inflammatory disorders and can even -in composite with DNA-trigger autoimmunity (43, 44), the anti-amyloidogenic activity of bacillaene could also contribute to prevent these additional negative effects of curli. Alternatively, purified bacillaene could also be envisioned for applications in combined therapies with other antibiotics to treat *E. coli* biofilm-associated infections. However, given the multiplicity of factors that contribute to antibiotic tolerance in biofilms, a general synergism between bacillaene and different classes of antibiotics is perhaps unlikely to be expected. Rather, based on its anti-curli potential, bacillaene is more likely to exhibit synergy specifically with those antibiotics that when used alone are tolerated by *E. coli* biofilms due to their binding by the network of curli fibers. Future studies exploring potential synergies between bacillaene and distinct antibiotics against curli-based *E. coli* biofilms will be helpful to increase knowledge on this aspect.

Lastly, and on a more general note, our findings also demonstrate the relevance of exploring microbial interactions beyond the exclusive search for antibiotics, as they can serve as platforms not only for finding compounds with novel and unique activities, but for uncovering additional functions of compounds previously characterized as antibiotics, as it is the case of bacillaene.

## MATERIALS AND METHODS

### Bacterial strains and growth conditions

*E. coli* W3110 is a reference K-12 strain from which derivate most *E. coli* strains used in this study. *E. coli* Δ*csgB* AR3110 is a derivative of W3110 in which pEtN-cellulose synthesis was restored by replacing a stop codon in *bcsQ* by a sense codon (45) and in which the *csgB* gene was deleted; hence, this strain produces pEtN-cellulose only. Additional derivatives of W3110 used in this study are Δ*csgD* and Δ*csgBA* deletion mutants and strains harboring a single chromosomal copy of *csgB::lacZ*, *csgD::lacZ* or *rpoS::lacZ* reporter fusions, all of which were reported previously (46, 47). *E. coli* MC4100 is also a K-12 strain but of a lineage that differ from W3110. LSR12 is a *csgDEFG/csgBA* deletion mutant in *E. coli* C600 background previously reported by the Chapman’s group (28). DAEC 2787 is a Diffusely Adhering *E. coli* strain proficient in curli production characterized by Alexander Schmitd’s group (Institut für Infektiologie ZMBE, Münster, Germany). *B. subtilis* strains used in the study are the NCIB 3610 strain and derivative mutants deficient in Sfp (*sfp::mls*; DS3337) or deficient in the synthesis of bacillaene (*pksL::cat*; DS4085), plipastatin (*ppsC::tet*; DS4114) or surfactin (*srfAC::Tn10 spec*; DS1122) (25). For routine handling, the strains were grown in solid or liquid LB medium at 28 or 37°C.

### Plasmid construction and transformation

To express the curli structural operon independent from its natural promoter we fused a synthetic RpoS-dependent promoter (SynP8(σ^s^) (48) to the *csgBAC* coding region on a low-copy number vector. In a first step, the pGBK27 vector containing SynpP8::*SFgfp* fusion was digested with *Eco*RI and *Xba*I enzymes to remove the *SFgfp* gene. Then, the *csgBAC* coding region was amplified by PCR using primers F_*Eco*RI_*csgBAC* (5’-GCCGGAATTCATGAAAAACAAATTGTTATTTATGA -3’) and R_*Xba*I_*csgBAC* (5’-ACTAGTCTAGATTAAGACTTTTCTGAAGAGGGC -3’) and genomic DNA from W3110 as template. Following *Eco*RI/*Xba*I digestion and purification, the PCR product was cloned into the *Eco*RI/*Xba*I-digested pGBK27 vector resulting in the pEC15 (pSynP(σ^s^)::*csgBAC*) plasmid. The correct ligation was confirmed by sequencing. Plasmid pEC15 was transformed into W3110 Δ*csgD* and Δ*csgBA* deletion mutants by electroporation with transformants being selected on LB agar supplemented with 100 μg/ml ampicillin.

To express and extracellularly export CsgA, *E. coli* LSR12 (Δ*csgDEFG/csgBAC*) was co-transformed by electroporation with plasmids pMC3 and pMC1, which encode his_6_-tagged CsgA and CsgG, respectively. Co-transformants were selected on LB agar plates containing 100 μg/ml ampicillin and 20 μg/ml chloramphenicol.

### Growth and imaging of macrocolony biofilms

Isolated or interacting bacterial macrocolonies were set on salt-free LB agar (49). When indicated, the agar was supplemented with ECM dyes (40 μg/ml Congo Red (CR), 20 μg/ml Coomassie brilliant blue (CBB) or 40 μg/ml Thioflavine (TS)), 40 μg/ml 5-bromo-4-chloro-3-indolyl β-D-galactopyranoside (X-gal) or different concentrations/volume of purified bacillaene or *pksL* extract. Interacting *E. coli-B. subtilis* macrocolonies were set by inoculating 5 μl of overnight cultures of each strain onto the agar with a separation distance of 1.5 cm. Macrocolonies were grown at 28°C for 3 days and imaged with a D3100 reflex camera (Nikon) equipped with a 12 mm extension tube for macro photography.

### Western blot and Plug-western blot analysis

Determination of CsgD cellular levels in *E. coli* macrocolonies was performed by western blot as previously described (27). Biomass from indicated macrocolony zones was collected, resuspended in phosphate-buffered saline (PBS), normalized by OD_578_ and pelleted down. SDS-PAGE was carried out using 12% acrylamide resolving gels and blotted onto polyvinylidene difluoride (PVDF) membranes. 7.5 μg cellular protein was applied per lane. A polyclonal serum against CsgD (custom-made by Pineda-Antikörper-Service, Germany), goat anti-rabbit IgG alkaline phosphatase conjugate (Sigma) and a chromogenic substrate (BCIP/NBT; PanReac AppliChem) were used for detecting CsgD proteins. Densitometric quantification of CsgD levels in the blots was performed using Fiji software.

Detection of the curli subunit CsgA in macrocolonies exclusively or in macrocolonies including the underlying agar (Plugs) was performed as described previously (27), with modifications. For macrocolony-only analysis, biomass of *E. coli* W3110 macrocolonies was collected at zones of interaction and distant (control) from *B. subtilis* 3610 wt or *pksL*, resuspended in PBS, normalized by OD_578_ and pelleted down. The pellets were treated with 5% SDS to efficiently depolymerizes the fibers. For plug analysis, *E. coli* W3110 macrocolonies were grown on agar poured in small wells with or without supplementation of bacillaene or extract from the *pksL* mutant. Subsequently, each colony including the underlying piece of agar (Plugs) was collected and treated with 5% SDS. For both macrocolony-only and plug samples SDS-PAGE were carried out in 15% acrylamide resolving gels and blotted onto 0.2 mm nitrocellulose transfer membranes. To control for potentially uneven sample application onto SDS gels, replicate gels were run in parallel and stained by CBB. A polyclonal serum against CsgA (kindly provided by Lynette Cegelski, Stanford University, US), goat anti-rabbit IgG alkaline phosphatase conjugate (Sigma) and chromogenic BCIP/NBT were used for detecting CsgA protein.

### *In vitro* CsgA polymerization

Freshly purified soluble CsgA was mixed in Eppendorf tubes with different concentrations of bacillaene or volume of *pksL* extract in the presence or absence of CR and diluted to a final concentration of 40 μM in KPi buffer. CsgA in KPi buffer and KPi buffer alone were established as positive and negative controls, respectively. The tubes containing the samples were incubated at 25°C for 16 h to allow CsgA polymerization. The presence or absence of white or red (in the presence CR) precipitate, indicative of the presence or absence of CsgA fibers, was visually recorded with a D3100 reflex camera.

To analyze the effect of bacillaene or *pksL* extract on the kinetics of CsgA polymerization, freshly purified soluble CsgA was mixed with different concentrations of bacillaene or volume of *pksL* extract and diluted to a final concentration of 20 μM in KPi buffer in wells of flat-bottom 96-well plates (Greiner Bio-One). Microtiter plates were covered with adhesive sealing film and the OD_578_ in the wells was measured for 16 h at 25°C, with readings every 20 min, on the Synergy 2 microplate reader (BioTeK), to monitor the kinetics of amyloid formation. The plate was shaken linearly for 5 sec to mix samples prior to each reading. All assays were performed in triplicates with at least three biological replicates. Representative OD_578_ curves for each condition are shown.

Additional information on experimental procedures is provided in Supplemental Materials and Methods.

## ACKNOWLEDGMENTS

We thank Ákos Kovács (DTU, Denmark) and Daniel Kearns (Indiana University, US) for the *B. subtilis* mutant strains. We also thank Paula Ruffatto (IBR) for technical assistance with bacillaene purification by HPLC. This study was supported by the Agencia I+D+i and ASaCTeI (Santa Fe) from Argentina (under grants PICT-2020-SERIEA-00534 and PEICID-2021-045 to D.O.S.) and Alexander von Humboldt Foundation from Germany (under grant Ref 3.4 – 8151/20 005 to D.O.S.). E.C is the recipient of a postdoctoral fellowship from National Research Council of Argentina (CONICET). M.I.Z, D.M.M. and D.O.S. are career investigators from CONICET.

## AUTHOR CONTRIBUTIONS

D.O.S. conceived the study and designed the experimental plan; most of the experiments were performed by E.C. with contributions from D.O.S.; M.I.Z. contributed to bacillaene purification; D.M.M. performed CsgB/A-bacillaene docking analysis; data were analyzed and interpreted by E.C. and D.O.S.; the paper was written by D.O.S. with inputs from the other co-authors.

We declare no competing interests.

## LEGENDS OF SUPPLEMENTAL FIGURES

**Fig. S1. (A)** Graph showing the number of *E. coli* viable cells at the zone of interaction or at a zone distant from *B. subtilis.* Cell biomass from each zone -as boxed in the image presented above the graph- was collected, resuspended and adjusted to the same OD_578_ and subjected to viable cell counting. **(B)** Macrocolony biofilms of *E. coli* AR3110 Δ*csgB* grown on CR-containing salt-free LB agar in close interaction with a macrocolony of *B. subtilis* 3610. The Δ*csgB* AR3110 macrocolony exhibits small intertwined wrinkles and a CR-based pink coloration characteristic of pEtN-cellulose. The images reveal that *B. subtilis* 3610 does not affect the morphological and CR-staining patterns of the Δ*csgB* AR3110 macrocolony. **(C)** Macrocolony biofilms of *E. coli* MC4100 and Diffusely Adhering *E. coli* (DAEC) 2787 grown on CR-containing salt-free LB agar in close interaction with a macrocolony of *B. subtilis* 3610. Images show that the inhibitory effect of *B. subtilis* on curli is reproduced in macrocolonies of other *E. coli* strains.

**Fig. S2.** Effect of Sfp-dependent secondary metabolites of *B. subtilis* 3610 on curli production and colony morphology examined at the microscale. **(A-E)** Macrocolony biofilms of *E. coli* W3110 grown on TS-containing salt-free LB agar in interaction with *B. subtilis* 3610 wt or derivative strains deficient in the phosphopantetheinyl transferase (*sfp*), surfactin (*srfAC*), plipastatin (*ppsC*) or bacillaene (*pksL*) (Left-hand side images). Fluorescence and merged fluorescence/phase contrast images of representative cross-sections through the W3110 macrocolony at zones of interaction or distant from respective *B. subtilis* strains (Right-hand side images). The spectral plots depict quantified patterns of TS fluorescence in respective cross-sections. Data provides further visual details demonstrating that in the absence of bacillaene only, curli production and hence the curli-dependent colony morphology occur normally.

**Fig. S3.** Bacillaene purification. **(A)** HPLC chromatograms at 362 nm showing bacillaene peaks in the processed *B. subtilis* wt extract, absent in the metabolite extract from the *pksL* mutant. **(B)** Absorption UV spectra of purified bacillaene fraction **(B)** and of metabolite extract from the *pksL* mutant **(C)**. The UV spectrum in B shows the characteristic band pattern of bacillaene, with absorption peaks appearing at 343, 362 and 383 nm, as previously reported (21). Consistent with this, bacillaene absorption bands are absent in spectrum shown in **C**.

**Fig. S4.** Lowest energy position of bacillaene on CsgA determined by docking simulations. **(A)** Representation of CsgA exhibiting binding of bacillaene to the surface of the β-helix. CsgA is shown with colour coding according to the type of residues: nonpolar residues (white), basic residues (blue), acidic residues (red), and polar residues (green). **(B)** Representation of CsgA β-sheet structure interacting with bacillaene.

**Mov. S1.** Time course of *E. coli* W3110 and *B. subtilis* 3610 macrocolonies growing in close interaction on CR-containing salt-free LB agar. Representative time-lapse movie that displays with good time resolution the inhibitory effect of *B. subtilis* on curli production in the *E. coli* macrocolony.

